# IFITM proteins inhibit HIV-1 protein synthesis

**DOI:** 10.1101/201525

**Authors:** Wing-Yiu Jason Lee, Chen Liang, Richard D Sloan

## Abstract

Interferon induced transmembrane proteins (IFITMs) inhibit the cellular entry of a broad range of viruses, but it has been suspected that for HIV-1 IFITMs may also inhibit a post-integration replicative step. We show that IFITM expression reduces HIV-1 viral protein synthesis by preferentially excluding viral mRNA transcripts from translation and thereby restricts viral production. Codon-optimization of proviral DNA rescues viral translation, implying that IFITM-mediated restriction requires recognition of viral RNA elements. In addition, we find that expression of the viral accessory protein Nef can help overcome the IFITM-mediated inhibition of virus production. Our studies identify a novel role for IFITMs in inhibiting HIV replication at the level of translation, but show that the effects can be overcome by the lentiviral protein Nef.

## Background

Viruses must replicate in cells; the consequence of such dependence is susceptibility. Cells have thus evolved a series of antiviral factors that directly subvert viral replication. Recent work has revealed the striking scale of antiviral mechanisms employed by cells - almost every step of viral replication is apparently targeted by host encoded antiviral factors. For HIV the best defined examples are APOBEC3G/3F and SamHD1 that inhibit viral reverse transcription, TRIM5α that targets viral capsid uncoating, and tetherin that prevents viral egress (reviewed in ^1,2^). However, genomic and proteomic screens can aid identification of further antiviral factors, and so there is an expanding list being described ^3,4^. Such studies identified the MxB/Mx2 protein that targets HIV-1 replication around nuclear entry ^5,6^, and the SERINC3 and SERINC5 proteins that target HIV-1 entry ^7,8^.

The interferon induced transmembrane proteins (IFITMs) have been similarly identified in multiple independent screens as antiviral factors ^9–12^. Humans have three antiviral IFITMs - IFITM1, IFITM2 and IFITM3 (reviewed in ^13^). IFITM expression inhibits viral entry into cells ^9,10^, a phenotype that is enhanced by incorporation of IFITM proteins into viral particles ^14–16^. Inhibition of viral entry is thought to occur via IFITM-mediated changes in the physical characteristics of the host cell membrane thereby inhibiting virus-cell membrane fusion ^17^, though other inhibitory mechanisms affecting entry have been proposed ^18^. By employing this generic inhibitory mechanism, IFITMs can then inhibit the entry of a broad range of viruses including influenza A virus, hepatitis C virus, Ebola virus, SARs coronavirus, Dengue virus, Zika virus, and HIV-1 ^9,10,19–21^.

Yet since the discovery of their antiviral effect upon HIV-1, it was immediately apparent that a viral replication step downstream of host genome integration might also be inhibited by IFITM proteins, as IFITM1 knockdown increased HIV-1 titres from infected CD4^+^ T-cells ^10^. In the same study, IFITM1 inhibited the replication of HIV-1 in T-cell culture despite the evidence that IFITM1 had no effect on viral replication steps from cell entry to host genome integration for the viral strain used. Other studies have also noted IFITM1-3 expression reduces viral particle production from cells ^12,14–16,22–26^. When this occurs, IFITM expression is typically associated with reductions in HIV-1 Gag levels, implying a block in viral protein synthesis.

Despite the frequency of these observations, there has been little to no attempt to explain them. Why viral production should be affected by IFITM expression is unclear. We therefore sought to investigate the potential block in protein synthesis induced by IFITMs and now show that IFITM expression leads to an inhibition of lentiviral production, which for HIV-1 occurs due to specific exclusion of viral mRNA from polysomes. Yet we also find that expression of the lentiviral accessory protein Nef can help relieve the inhibition in virus production, allowing HIV-1 to better replicate in the presence of IFITM proteins. Thus IFITM proteins are able to inhibit HIV-1 replication through three distinct processes - inhibition of viral entry, reduction of viral particle infectivity, and inhibition of viral protein synthesis.

## Results

### IFITMs inhibit HIV-1 production

We first wanted to confirm if IFITMs affect the production of HIV from virus producing cells as noted in previous reports ^10,12,14–16,22–26^. We used a co-transfection scheme in which we co-transfected HEK293T cells with plasmids containing HIV-1 proviral DNAs together with expression vectors encoding human IFITM1-3, allowing the effects on virus production to be studied independently of effects of IFITM proteins on viral entry. The quantity of virus produced was then analyzed by supernatant p24 ELISA and reverse transcriptase activity assays. Increasing levels of IFITM expression decreased HIV-1 NL4-3 output by in a concentration-dependent manner consistent with loss of intracellular p55 and p24 levels (Figure 1A, 1B), with a similar decrease seen in supernatant when both p24 concentration (Figure 1C) and reverse transcriptase activity (Figure 1D) were measured. Notably, the expression of IFITM1 and IFITM2 reduced the HIV-1 output while the effect was weaker for IFITM3. We then studied the effect of IFITMs on other viral proteins, and found IFITM1 and IFITM2 expression led to a reduction in the levels of all HIV-1 NL4-3 proteins assayed in transfected cells (Figure 1E). Similar to HIV-1 NL4-3, viral production from the HIV-1 plasmid proviral DNAs 89.6 and Indie-C1 was also reduced by IFITM expression (Figure 1C and D).

In addition to HIV-1, the expression of IFITMs also reduced the quantity of virus produced from cells transfected with plasmid proviral DNAs for HIV-2 and SIVs (Figure S1A and S1B). Of note, HIV-2_ROD_ and SIV_AGM_ appeared to be less susceptible to this IFITM-mediated effect compared to the HIV-1 viruses studied. We also found similar inhibitory effects upon HIV-1 with transfected African Green Monkey IFITM1 (Figure S1C).

**Figure 1.**
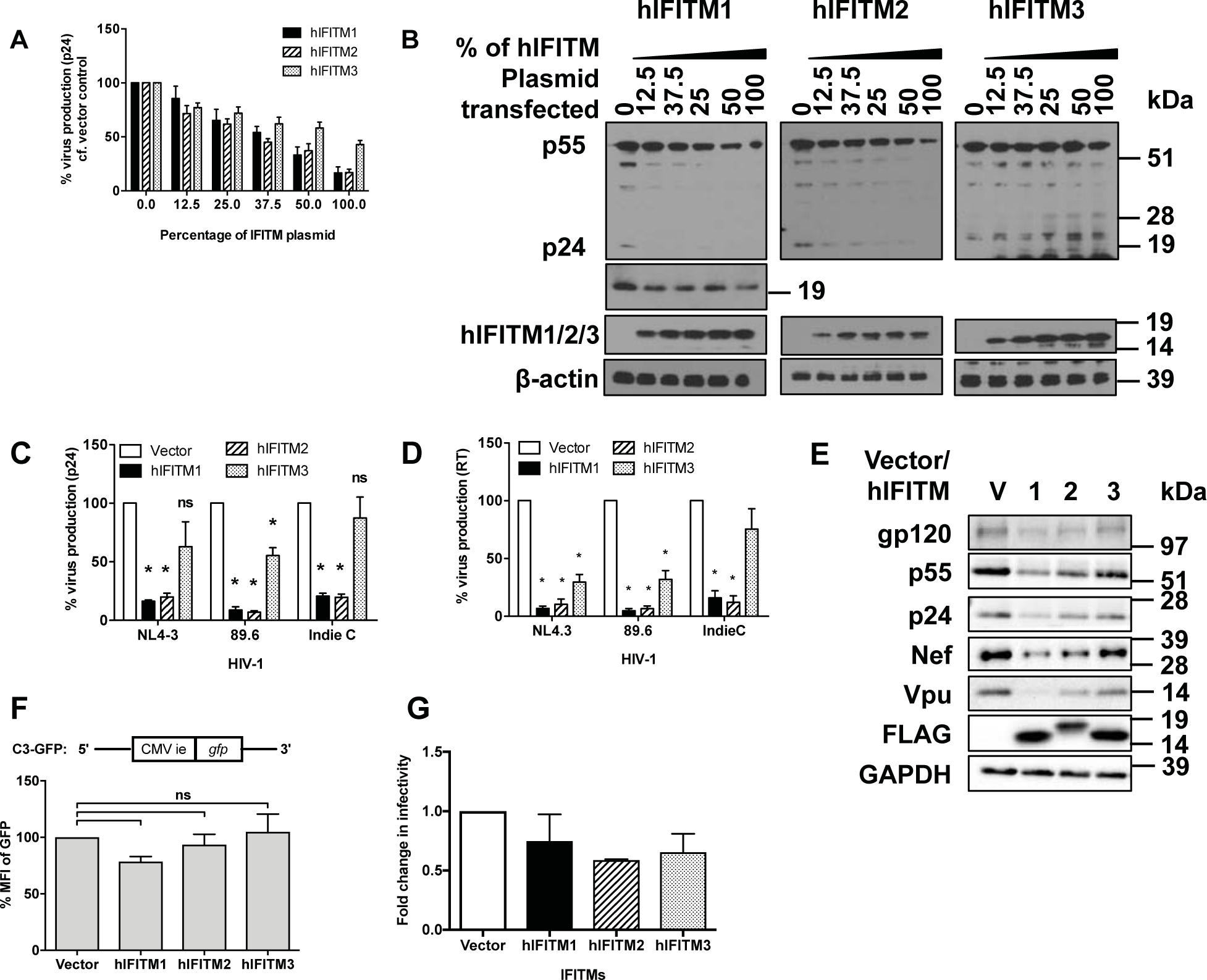
IFITMs inhibit HIV viral output and viral protein production in transfected cells. HEK293T cells were transfected with a titration of expression vectors for FLAG tagged IFITMs. **(A)** Levels of virus production were measured by p24 ELISA 48h post-transfection, and **(B)** cellular viral proteins and IFITMs were analyzed by immunoblotting. HEK293T cells were transfected with 0.5μg of FLAG-tagged IFITM expression vectors and 0.5μg of HIV-1 NL4-3 proviral DNA. Levels of virus production from indicated HIV-1 proviral DNAs were measured by **(C)** p24 ELISA and **(D)** reverse transcriptase (RT) activity assay and **(E)** immunoblotting of intracellular proteins 48h post-transfection. **(F)** Level of GFP expression in HEK293T cells co-transfected with 0.5μg CMV-driven GFP vector and 0.5μg IFITM-expression vectors or empty vectors measured by flow cytometry 48 hours post-transfection. **(G)** HIV-1 NL4-3 virus was produced from HEK293T cells transfected with vector or the indicated IFITMs in TZM-bl cells for 48 hours. Infectivity in TZM-bl cells was measured by luciferase activity assay 48 hours after infection with virus stock with equivalent p24 concentration. Data show mean + S.E.M. of more than 3 independent experiments. All differences were assessed with Student’s t-test and * indicates p<0.05.

We were concerned however that the loss of viral production we saw might be due to generic inhibition of protein production during IFITM expression, but detection of GFP from a transfected reporter plasmid was not significantly affected by IFITM expression (Figure 1F). We also measured the infectivity of the virus produced from IFITM-expressing cells, in TZM-bl cells and showed that virus produced have lower infectivity (Figure 1G), in agreement with previous results ^14–16^.

We noted that levels of IFITM proteins in transfected 293T cells were often higher than those seen in type-I interferon treated CD4+ T-cells and monocyte derived macrophages (MDMs) (Figure S2) and so we next investigated the influence of IFITM proteins in cells in which levels of expression could be inducibly expressed. While we also wanted to test if IFITMs inhibit viral production in infected cells rather than provirus transfected cells. As such we infected SupT1 cells expressing doxycycline inducible IFITM1-3 with wild type NL4-3 virus ^10^. To avoid the well documented inhibition of virus entry by IFITMs playing a role in our experiments, after spinoculation, unbound virus was washed off and cells were resuspended in new medium containing doxycycline and the CXCR4 antagonist AMD3100 to induce IFITM expression only after virus entry and to limit viral replication to a single round respectively. Complete entry blockade was confirmed with pre-treatment of cells (Figure S2A). The quantity of wild type HIV-1 produced by IFITM-expressing T-cells was inhibited across a range of viral inputs and was paired with losses in all viral protein levels, but with p55 levels somewhat less affected (Figure 2B). Though collectively this indicates that IFITMs inhibit viral production independent of their effects on viral entry. Notably, the levels of doxycycline induced IFITM were similar to those found in type I interferon treated CD4+ T-cells and monocyte derived macrophages (Figure S2B, C and E).

**Figure 2.**
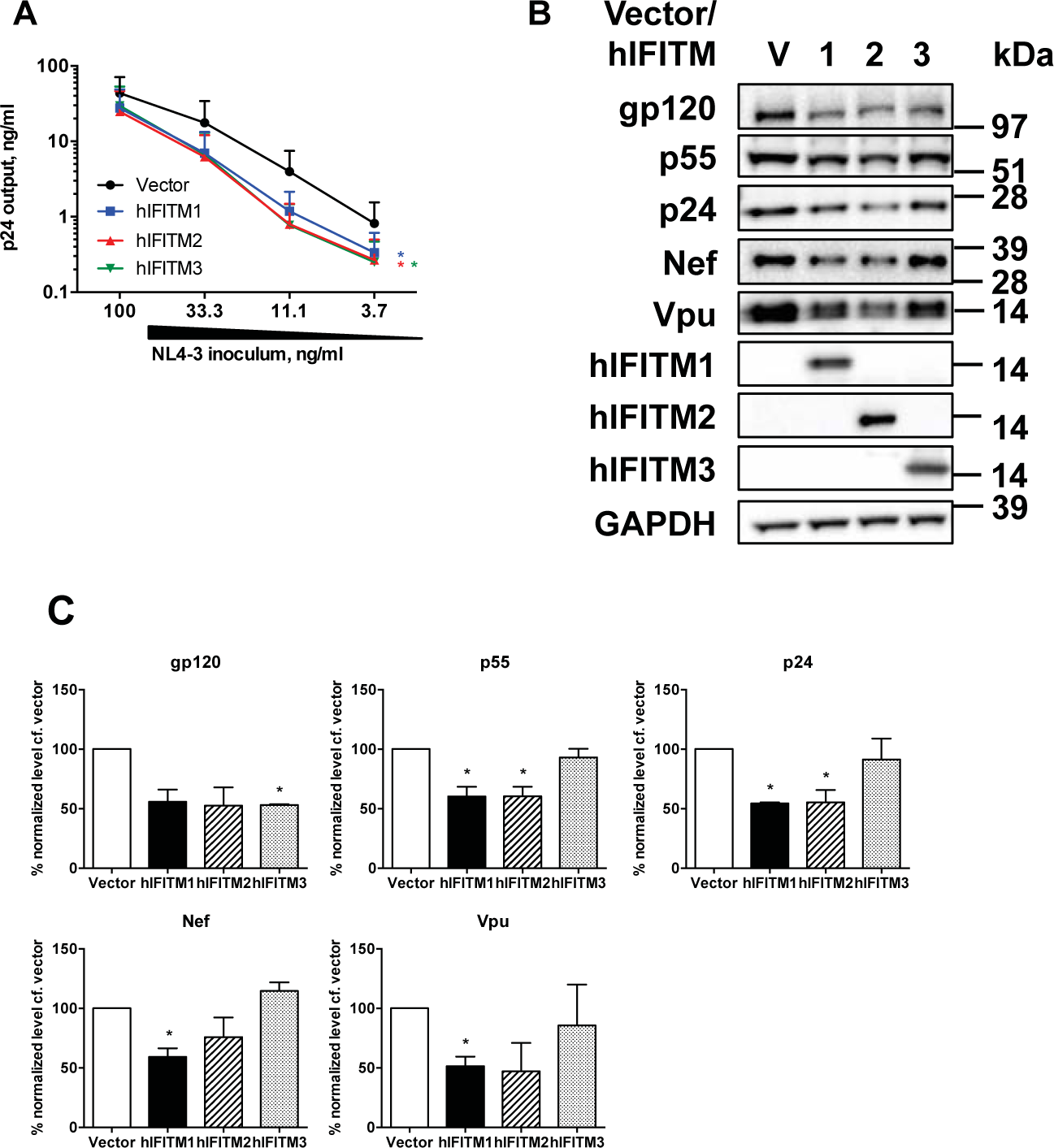
Inducible expression of IFITMs after viral entry inhibits HIV viral output and viral protein production in infected SupT1 cells. **(A)** SupT1 cells were infected with the indicated dilutions of wild type HIV-1 NL4-3 inoculum and then treated with 1μg/ml doxycycline to induce IFITMs post-entry and 5μM AMD3100 to limit infection to a single cycle. At 72h post-infection levels of virus production were measured by p24 ELISA and differences assessed by Two-way ANOVA with Bonferroni’s multiple comparison test. **(B)** Cellular viral proteins and IFITMs were analyzed by immunoblotting and **(C)** densitometry. Data show mean + S.E.M. of 3 independent experiments. Differences were assessed with Student’s t-test and * indicates p<0.05.

We next wished to investigate this effect using knockdown of endogenous IFITM levels to avoid overexpression artefact, and to assess the relative contribution of IFITMs towards inhibition of viral production that is observed with type I interferon (IFN) treatment. We therefore analyzed viral output from HIV-1 NL4-3 plasmid proviral DNA transfected and IFN-β treated TZM-bl cells that had been transduced with shRNAs targeting different IFITMs ^27^. Infection of cells with virus produced during transfection was inhibited through the use of the CXCR4 antagonist AMD3100, and thus the level of viral output in our analysis cannot be confounded by entry inhibition due to IFITM expression. The efficiency of knockdown in IFITM expression was confirmed by immunoblotting using monoclonal antibodies specific for individual IFITMs (Figure 3A). As anticipated, IFN-β treatment reduced viral production in HIV-1 transfected TZM-bl cells (Figure 3B). Despite some weak knockdown cross-reactivity, such that shRNA targeting IFITM1 caused weak knockdown of IFITM2 (Figure 3A), and that it was not possible to knockdown IFITM3 in isolation, knockdown of both IFITM1 and IFITM2 clearly rescued the level of virus produced in IFN-β treated cells (Figure 3C). Further, knockdown of all three IFITMs gave a higher rescue, indicating that the inhibitory effects of IFITM1 and IFITM2 on viral production are perhaps non-redundant in the context of HIV-1 replication. These effects were paired with rescue of all viral proteins levels during IFITM knockdown (Figure 3D and 3E). Knockdown of IFITMs rescued viral protein production by around 1.5 to 3.5 fold, and so is concordant with viral production rescue of 2.0 to 2.5-fold. These data then show that endogenous levels of IFITMs can inhibit viral particle output and that our prior results in 293T transfections and SupT1 infections are not an overexpression artefact. They also suggest that IFITMs are a significant component of the antiviral response induced by type I interferon.

**Figure 3.**
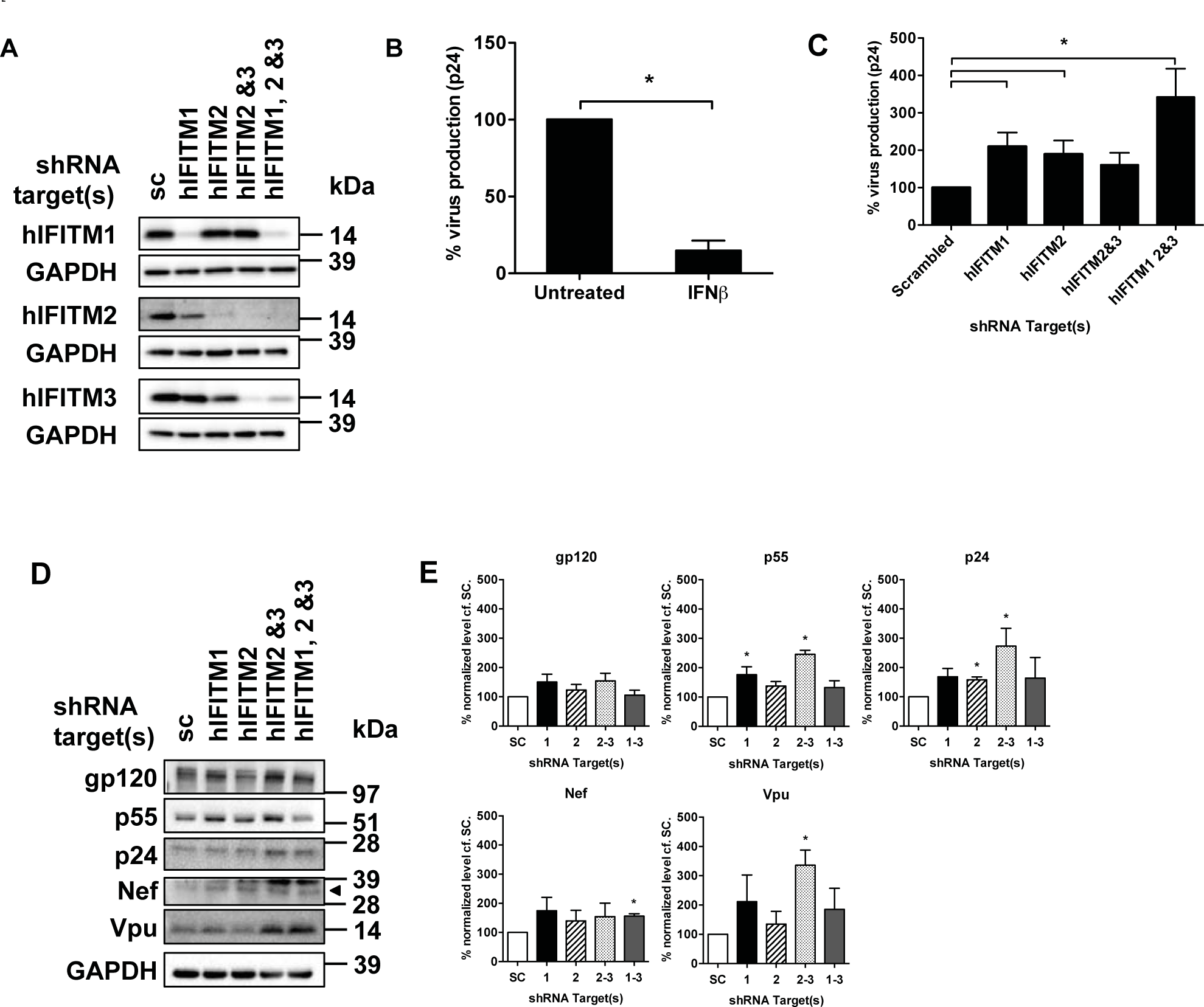
Knockdown of IFITMs rescues HIV-1 output and viral protein production in TZM-bl cells. TZM-bl cells transduced with shRNAs against the indicated IFITMs or scrambled control (sc) were treated with 100IU/ml IFNβ for 72 hours. **(A)** Levels of IFITM expression were analyzed by immunoblotting. **(B)** Scrambled control TZM-bl cells were transfected with HIV-1 NL4-3 proviral DNA and treated with 100IU/ml IFNβ. Level of virus production was measured by p24 ELISA 72h post-transfection. TZM-bl cells transduced with shRNAs against the indicated IFITMs or scrambled control were transfected with HIV-1 NL4-3 proviral DNA and then treated with AMD3100 and IFNβ 4h post-transfection. Level of virus output and viral proteins was measured by **(C)** p24 ELISA and **(D)** immunoblotting, respectively, at 72 hours post-transfection. **(E)** Immunoblotting was further analyzed by densitometry. Data show mean + S.E.M. of 3 independent experiments. Differences were assessed with Student’s t-test and * indicates p<0.05.

We next sought to confirm if the knockdown of physiologically relevant levels of IFITMs in HIV-1 infected primary cells could rescue viral protein production. Primary human CD4^+^ T cells were infected with HIV-1 following transduction with lentiviral particles for the expression of shRNAs targeting different IFITMs. Transduction with one shRNAs resulted in a specific ~90% downregulation of IFITM2, while another shRNA led to a ~50% downregulation of all three IFITMs (Figure 4A-C). We were unable to specifically knockdown IFITM1 with reasonable efficiency. To mitigate the effects of IFITMs inhibiting viral entry on our studies of inhibited viral production, we took advantage of recent observations showing that different IFITM proteins vary in their inhibition of viral entry depending on HIV-1 co-receptor usage ^16,22,28^. IFITM1 inhibits entry of CCR5-tropic viruses and IFITM2/3 predominantly inhibits CXCR4-tropic viruses. We therefore studied viral production effects driven by IFITM2 and IFITM3 with dual-tropic HIV-1 89.6, wherein cells were treated with the CXCR4 antagonist AMD3100 prior to and during infection. AMD3100 treatment would direct viral entry via CCR5 due to CXCR4 blockade and thus mitigate entry inhibition by either IFITM2 or IFITM3. We used intracellular p24 staining of infected cells in single cycle infections to measure uptake of virus after infection with trypsin treatment to remove cell surface bound virus ^29–31^. Although cellular uptake of viral p24 is not an exact measure of virus entry due to the uptake of virus into potentially non-productive endocytic pathways, p24 uptake is commonly found to be inhibited by IFITM expression ^14,15,24,25^, while endocytic uptake of virus occurs at a lower rate in primary CD4^+^ T-cells ^32^. We confirmed that viral uptake was minimally impeded with these during IFITM knockdown (Figure 4D and E), consistent with prior observations ^28^. Therefore reductions in virus production were considered to arise predominantly from IFITM-driven effects other than inhibition of viral entry. As such knockdown of IFITM2 led to a clear rescue of viral particle output (7-fold) that was coupled with rescue in viral protein expression (Figure 4F,G). Similarly, knockdown of IFITM1-3 led to rescue of viral protein production and viral particle output (>10-fold) (Figure 4F, G and H). Though as IFITM1 which was depleted in this experiment can inhibit R5-tropic virus entry, a proportion of this latter result for IFITM1-3 may be attributable to weak IFITM1 driven entry inhibition. That this effect was very weak (Figure 4F and G) suggests that viral production inhibition may be the dominant form of IFITM driven inhibition in this experiment. Overall, this confirms that physiologically relevant levels of IFITM proteins can clearly inhibit HIV-1 production in primary cells independent of their effects upon entry and infectivity, that this is not an overexpression artefact, and occurs in a manner that is consistent with our observations in cell lines.

**Figure 4.**
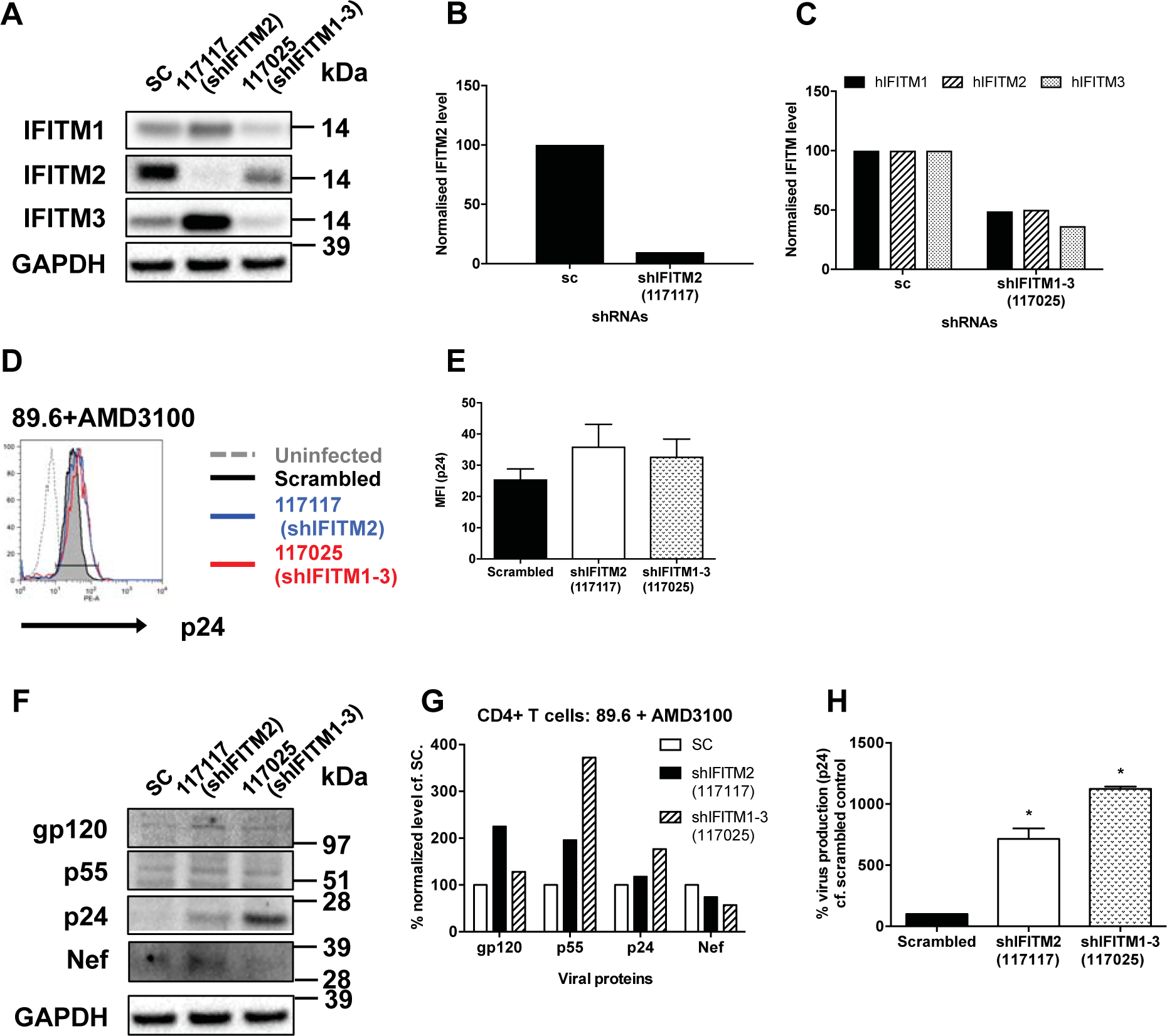
Knockdown of IFITMs rescues HIV-1 output and viral protein production in primary human T cells. Activated human CD4^+^ T cells were transduced with lentivirus expressing the indicated shRNAs against IFITMs or scrambled sequence (sc) for 48 hours. **(A)** Level of IFITM expression was analyzed by immunoblotting and (B) densitometry; data was normalized to GAPDH and scrambled control. Cells were then infected with HIV-1 89.6 with equivalent p24 concentrations. Cells were pre-treated with AMD3100 for 2 hours prior to infection and throughout. viral proteins in infected cells was analyzed by immunoblotting. Immediately after infection, cells were treated with trypsin and washed with PBS before intracellular staining of p24 and flow cytometry to measure virus uptake. Intracellular levels of p24 and median fluorescence intensity are shown in **(D)** representative histograms and **(E)** summary bar chart. Levels of **(F)** viral proteins in infected cells from one of the blood donors were shown by **(G)** densitometry. **(H)** Level of virus production was measured by p24 ELISA 72 hours post-infection and normalized. Data show mean + S.E.M. of 2 blood donors. Differences were assessed with Student’s t-test and * indicates p<0.05.

### IFITMs restrict HIV-1 protein synthesis

As we found that IFITM expression was associated with losses in viral protein production and viral output, we next asked if IFITMs play a role in inhibiting the transcription or stability of viral mRNA by analyzing the level of unspliced, singly-spliced, and multiply-spliced transcripts detected by RT-qPCR from total RNA of whole cell lysates. IFITM1 and IFITM2 weakly reduced the levels of singly-spliced and multiply-spliced transcripts, while IFITM3 had no measureable effect (Figure 5A). The total level of unspliced transcripts remained unaffected in the presence of IFITM proteins (Figure 5A) but nonetheless there were reductions of intracellular level of Gag (a product of unspliced transcripts) in our prior analyses (Figures 1–4). This prompted us to investigate if IFITM proteins affect translation of viral transcripts.

**Figure 5.**
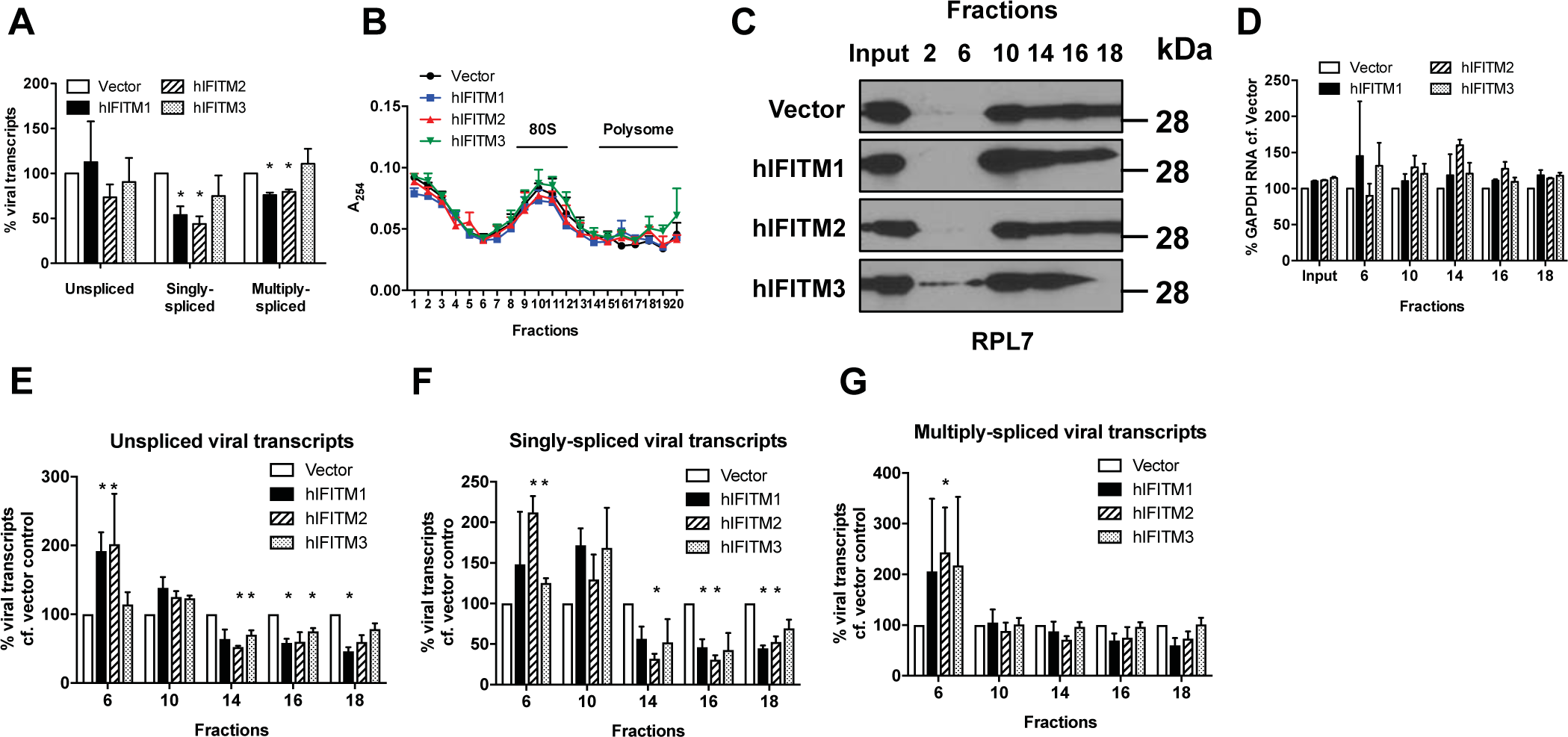
IFITM proteins inhibit HIV-1 protein synthesis. Activated human CD4^+^ T cells were transduced with lentivirus expressing the indicated shRNAs against IFITMs or scrambled sequence (sc) for 48 hours. **(A)** Level of IFITM expression was analyzed by immunoblotting and (B) densitometry; data was normalized to GAPDH and scrambled control. **(A)** Level of viral transcripts in HEK293T cells transfected with HIV-1 NL4-3 and IFITM DNA was measured by qPCR 48 hours post-transfection and the data normalized. **(B)** Level of RNA in sucrose gradient fractions of HEK293T cells transfected with proviral HIV-1 NL4-3 DNA and expression vectors for IFITMs (or vector alone) was measured by absorbance at 254nm. **(C)** Level of ribosomal protein L7 was analyzed by immunonblotting to identify fractions enriched for polysomes. **(D)** Level of GAPDH RNA in IFITM-expressing cells was normalized to vector control in the indicated sucrose gradient fractions. Level of **(E)** unspliced, **(F)** singly-spliced and **(G)** multiply-spliced viral transcripts in the indicated sucrose gradient fractions in **(B)** was analyzed by qPCR then normalized with the levels of GAPDH RNA and input RNA. Data shows mean + SEM of 3 independent experiments. Differences were assessed by Student’s t-test and * denotes p<0.05 compared to vector control.

To achieve this we performed a polysome analysis for HIV-1 transcripts during IFITM expression. We first analyzed the RNA content in cell lysates fractionated on a sucrose gradient by ultracentrifugation. The total RNA profiles (Figure 5B) RNA recovered from selected fractions of virus producing cells expressing IFITMs are comparable to vector-transfected control, suggesting that IFITM expression did not alter global translation in the cell. We used the level of ribosomal protein L7 as a guide for selection of fractions in our analysis (Figure 5C) as it is a component of 60S ribosomes, which are enriched in 80S monosomes (such as fraction 10 Figure 5C) as well as polysomes (such as fractions 14-18. The levels of GAPDH mRNA were not reduced in polysome fractions. Reductions in the levels of viral transcripts in polysome fractions (in contrast to whole cell lysates – Figure 5A) would indicate specific translational inhibition. As such, we analyzed the level of viral transcripts in selected fractions covering the whole RNA profile with RT-qPCR.). IFITM1 and IFITM2 significantly reduced the level of unspliced and singly-spliced viral transcripts in polysomes (Figure 5E and F) and increased the level of these transcripts in the ribosome-free fraction (fraction 6), suggesting that expression of IFITM1 and IFITM2 inhibited translation of these viral transcripts. IFITM1 and IFITM2 also appeared to somewhat reduce the level of multiply-spliced transcripts (Figure 5G), though the effect did not reach statistical significance, but was nonetheless coupled with a clear accumulation of ribosome-free multiply-spliced transcripts in low concentration sucrose fraction (Figure 5G - fraction 6) relative to vector control, similar to other viral transcript classes. These data were consistent with our analysis of viral production (Figure 1) wherein IFITM1 and IFITM2 expression lead to a more potent block in viral production than for IFITM3. Thus IFITM proteins inhibit translation of HIV-1 transcripts by leading to their specific exclusion from polysomes, resulting in reduced levels viral proteins and reduced viral output.

### HIV-1 RNA is a determinant of IFITM-mediated restriction of protein synthesis

To better understand how IFITM proteins help suppressing viral translation, we explored the possibility that they may be involved in a process that is able to distinguish viral RNA from cellular RNA. To address this, we measured the level of unspliced viral RNA transcripts in the polysome fraction of HEK293T cells transfected with a codon-optimized vector for HIV-1 NL4-3 Gag only ^33^ in conjunction with IFITM-expression vectors. Codon-optimization changes the codon bias of the construct towards and therefore also alters the secondary structures of viral RNA. Changing codon bias thus renders HIV-1 unspliced and singly-spliced transcript expression independent of the HIV Rev responsive element (RRE), a *cis* acting RNA structure necessary for transcript nuclear export ^34^. Codon optimization of *gag* rescued the level of unspliced HIV-1 transcripts in polysomes during IFITM expression, nor was there an accumulation of transcripts in low sucrose fractions as was seen with wild type NL4-3 (fraction 7 in Figure 6A compared to fractions 6 in Figure 5E, F and G). As a result, the level of extracellular p24 production from cells expressing IFITM1 and IFITM2 was not only rescued but enhanced, and the production of the *gag* products p55 and p24 was also restored (Figure 6B, C and F). Notably p24 protein production was only partially rescued, but p55 was entirely rescued (Figure 6F). Contrasting data for full length pNL4-3 DNA show clear losses of viral production and protein expression in similar experiments (Figure 6D, E and F).

**Figure 6.**
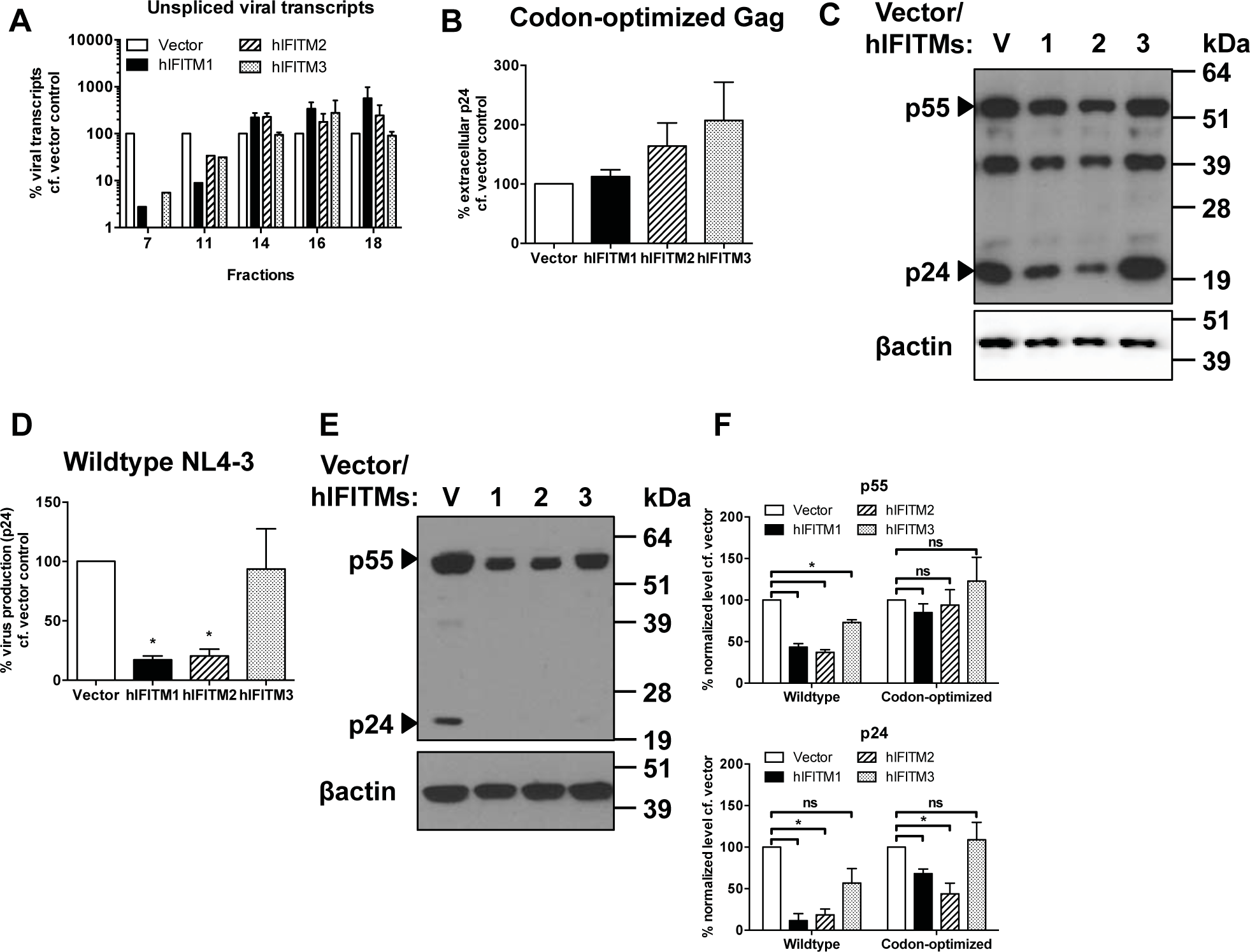
HIV-1 RNA is a determinant of IFITM-mediated inhibition of protein synthesis. **(A)** Normalized levels of unspliced viral transcripts (measured by qPCR) in the polysome fractions of HEK293T cells transfected with codon-optimized HIV-1 NL4-3 Gag DNA and IFITM expression vectors at 48h post-transfection. **(B)** Levels of extracellular p24 in HEK293T cells transfected with codon-optimized HIV-1 Gag DNA measured by p24 ELISA 48h post-transfection and normalized. Cellular levels of HIV-1 Gag (p55 and p24) were analyzed by **(C)** immunoblotting. **(D)** Levels of virus production in HEK293T cells transfected with 0.5μg wild-type HIV-1 NL4-3 proviral DNA and 0.5μg IFITM-expression plasmids or empty vector was measured by p24 ELISA 48 hours post-transfection and normalized. Cellular levels of HIV-1 Gag (p55 and p24) in cells transfected were analyzed by **(E)** immunoblotting and **(F)** densitometry. Data show mean + SEM of 3 independent experiments and differences were assessed by Student’s t-test, * denotes p<0.05 compared to vector control.

We then sought to identify specific RNA features that may render HIV-1 susceptible to an IFITM-driven translational block. As our findings of transfected HEK293T cells showed that unspliced and singly-spliced viral transcripts are more readily excluded from polysomes by IFITM proteins than multiply-spliced transcripts in those cells (Figure 5), we reasoned that RRE, which is present in only unspliced and singly-spliced transcripts may be somehow recognized during IFITM protein expression leading to translational inhibition. To address this, we exploited constructs in which singly-spliced and unspliced transcripts had been rendered RRE and Rev independent via the introduction of the Mason Pfizer Monkey Virus (MPMV) constitutive transport element (CTE) ^35^. A modification which allows all transcripts to be exported from the nucleus by the cellular TAP/NXT1 pathway, rather than RRE bearing Rev-dependent transcripts exporting via the CRM1 pathway ^36^. We found that CTE dependent Rev independent virus was still inhibited by IFITM proteins (Figure S3), demonstrating at least that nuclear export pathway does not affect the virus production inhibition seen with IFITM proteins. Further introduction of the RRE into CTE bearing constructs did also not augment the restriction.

Overall, these data suggest that the restriction of translation by IFITMs occurs at the RNA level, though it is not clear that the RRE plays a significant role in susceptibility, despite that RRE bearing transcripts are more affected in translation. Therefore some other distinguishing feature of viral transcripts or translation determines their susceptibility to IFITM proteins.

### HIV Nef can help overcome IFITM-mediated restriction of virus production

Under the selective pressure of cellular restriction factors, retroviruses have evolved to counteract surveillance and restriction by host antiviral factors (reviewed in ^1^). We therefore sought to investigate if HIV is able to counteract the restriction of viral production by IFITM proteins. We reasoned that HIV-1 Nef may influence this process. IFITMs are membrane proteins and lentiviral Nef regulates the trafficking of many membrane proteins such as CD4, MHC-I and tetherin to aid immune evasion and viral replication ^37–39^. Further, the expression of a dominant negative form of HIV-1 Nef ^40^, or *nef* deletion has previously been linked to deficits in viral production ^41^.

We therefore measured the level of virus production in HEK293T cells transfected with expression vectors of IFITMs with wild type or *nef-*deleted HIV-1 NL4-3 proviral DNA. The level of *nef*-deleted virus produced was inhibited 18-fold by IFITM1, 21-fold by IFITM2 and 5-fold for IFITM3, suggesting a 4-fold enhancement of virus production due to HIV-1 Nef during IFITM1 and IFITM2 expression (Figure 7A). We confirmed a significant reduction in HIV-1 Gag (p55) levels in cells producing *nef*-deleted virus while IFITMs are expressed (Figure 7B, S4B), indicating that Nef may rescue the translational suppression driven by IFITMs. As these data were generated through transfection, they were independent of any influence of Nef upon viral entry ^7,8^. It is also of note that exogenous IFITM levels were not reduced by HIV-1 Nef, perhaps indicating that antagonism of IFITM proteins may not occur via degradation (Figure 7B).

**Figure 7.**
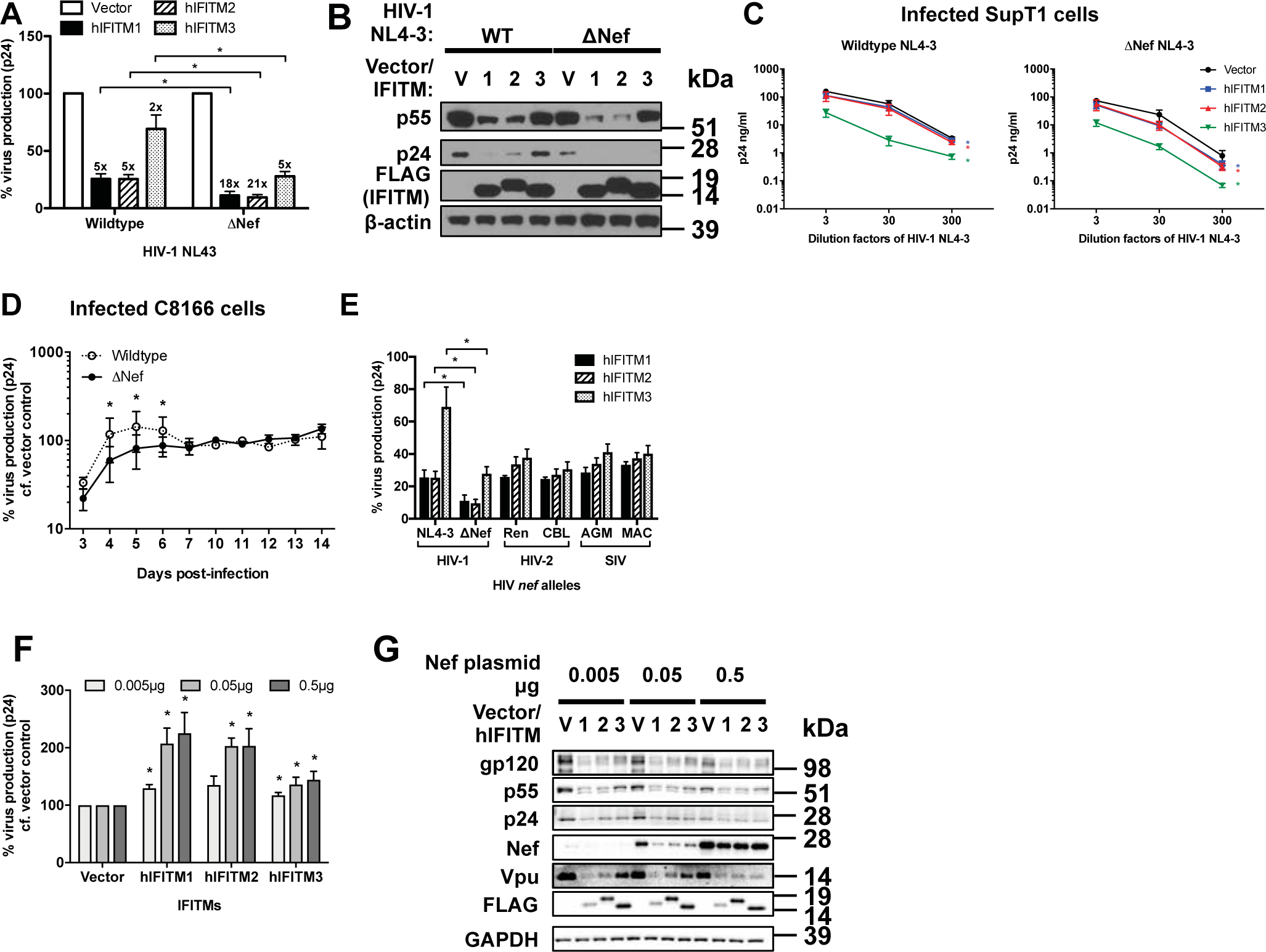
HIV Nef can help overcome IFITM-mediated restriction of protein synthesis. **(A)** Level of virus production in HEK293T cells transfected with 0.5μg expression vectors for IFITMs and 0.5μg HIV-1 NL4-3 proviral DNA with a deletion (ΔNef) was measured by p24 ELISA 48 hours post-transfection and the data normalized, fold change of virus production compared to vector control is indicated. Differences were assessed with Student’s t tests. **(B)** Intracellular level of p55/p24 and IFITMs was measured by immunoblotting. **(C)** SupT1 cells were infected with the indicated dilutions of wild type or Nef-deleted (ΔNef) HIV-1 NL4-3 inoculum and then treated with 1μg/ml doxycycline to induce IFITM expression post-entry and 5μM AMD3100 to limit infections to a single round. Level of virus production was measured by p24 ELISA 72 hours post-infection. Differences were assessed with Two-way ANOVA and Bonferroni post-tests. **(D)** C8166 cells constitutively expressing either vector control or IFITM1 were infected with either wild type or Nef-deleted (ΔNef) HIV-1 NL4-3. Levels of virus production were measured by p24 ELISA at the indicated time-points post-infection and were normalized to the levels of virus produced from vector controls. Differences were assessed with Two-way ANOVA. **(E)** HEK293T cells were transfected with 0.5μg expression vectors for IFITMs and 0.5μg HIV-1 NL4-3 proviral DNA with either wildtype NL4-3 *nef* or the indicated lentiviral *nef* alleles. Virus production was measured by p24 ELISA 48 hours post-transfection and the data normalized. Differences were assessed with Student’s t tests. **(F)** HEK293T cells were transfected with ΔNef HIV-1 NL4-3 proviral DNA, expression vectors for IFITMs and an increasing proportion of HIV-1 Nef-encoding vector versus empty vector in a fixed total quantity of 1μg. Level of virus production was measured by p24 ELISA 48 hours post-transfection, while levels of viral proteins and IFITM-FLAG expression was analyzed by **(G)** immunoblotting. Differences were assessed by Student’s t-test. All data show mean + SEM from 3 independent experiments and * denotes p<0.05.

We next investigated if Nef presented an advantage to viral growth in infected T-cells. We infected SupT1 cells with either wild type or *nef*-deleted NL4-3 virus by post-entry induction of IFITM expression via doxycycline (Figure 7C). As *nef*-deleted virus has an inherent growth defect, we normalized the quantity of wild type and *nef*-deleted virus produced from IFITM-induced cells to vector control cells. We chose to induce IFITMs after X4-tropic HIV-1 NL4-3 entry and limit infections to a single round through the use of the CXCR4 antagonist AMD3100 after infection to mitigate the influence of IFITMs upon viral entry. In this regard, we found that *nef-*deleted virus was no more susceptible to IFITM-mediated restriction of entry than wild type virus (Figure S4A). Compared to wild type virus, production of *nef*-deleted virus in infected SupT1 cells was reduced significantly during IFITM expression, surprisingly IFITM3 showed the greatest degree of rescue in SupT1 cells (Figure 7C), in contrast to data for transfected 293T cells (Figure 7A and B).

To further confirm the role of IFITM1 in suppressing virus production in T cells, we monitored the production of either wild type or *nef*-deleted HIV-1 NL4-3 virus in multiple cycle infections in a C8166 T-cell line transduced to constitutively over-express IFITM1. As IFITM1 does not typically affect the entry of CXCR4-tropic virus such as NL4-3 (Figure 4A, B and ^10,28^), any difference in replication measured between wild type and *nef*-deleted virus could only arise from IFITM1-mediated effects on viral production, not cell entry. Therefore this experiment was not pursued for IFITM2 and IFITM3, as their constitutive expression would inhibit NL4-3 virus entry and complicate interpretation. We found that in the presence of IFITM1, the growth of *nef*-deleted virus was significantly reduced relative to wild type virus, with wild type virus showing 2-fold inhibition at 5 days post-infection, while *nef*-deleted virus was inhibited 5-fold by IFITM1 (Figure 7D). As wild type and *nef*-deleted virus data in C8166 cells were individually normalized to the same viruses in C8166 vector control cells, the differences observed are not likely due to Nef overcoming SERINC3/5 expression ^7,8^, as these factors would be similarly present in both cell lines. Interestingly, the benefit of Nef expression in the context of IFITM1 was lost in the later stages of viral culture, when cell-to-cell infection typically predominates ^42^. As *nef*-deleted virus was no more susceptible to IFITM-mediated restriction of entry than wild type virus (Figure S4A), these findings are only due to the influence of Nef to partially rescue virus production as seen in 293T cell transfections and SupT1 infections (Figure 7A-C).

We then investigated if Nef from other lentiviruses was able to overcome IFITM-mediated inhibition of protein synthesis using an HIV-1 NL4-3 plasmid proviral DNA into which lentiviral *nef* alleles had been substituted ^43^. The level of virus production for HIV-1 NL4-3 bearing other *nef* alleles during IFITM expression was generally comparable to levels for wild type NL4-3 *nef* (Figure 7E), indicating that ability of lentiviral Nef proteins to overcome IFITM mediated restriction of virus production is common.

To further confirm the role of Nef in antagonizing IFITMs, we titrated a plasmid bearing NA7 *nef* into cells transfected with IFITM expression plasmids and *nef*-deleted NL4-3 proviral HIV-1 DNA. Nef expression *in trans* rescued viral production from IFITM-mediated inhibition in a dose dependent manner (Figure 7F). However, we note overexpression of Nef in cells transfected with 0.5μg Nef-encoding plasmid resulted in a overall lower level of viral proteins compared with cells expressing lower levels of Nef. Interestingly, increased Nef levels helped restore viral production in the context of IFITM expression (Figure 7F), this rescue was not reflected in viral protein production levels (Figure 7G and S4C), implying that the ability of Nef to rescue HIV production during IFITM expression does not occur at the level of protein synthesis. In summary, these data show that Nef can help overcome IFITM-mediated inhibition of HIV-1 production in a concentration-dependent manner. Taken together, our data demonstrate that HIV-1 Nef is able to help overcome the inhibition of viral protein production exerted by IFITM proteins and therefore enhance infection.

## Discussion

We report here that IFITM protein restrict HIV-1 by excluding viral mRNA from polysomes thereby specifically inhibiting protein synthesis. By variously using transfection, post-entry IFITM induction, and viral co-receptor bypass strategies, we have identified a viral entry independent effect due to IFITMs that inhibits viral production. Since the discovery of the antiviral effect of IFITM proteins upon HIV-1 entry and infectivity, there have been data suggesting that there may be a distinct IFITM function directed towards HIV-1 production ^10,12,14,15,22–26^. Despite the relative frequency of these passing observations on viral production, the underlying processes leading to loss of viral production during IFITM expression have not been explored.

We have demonstrated the breadth of the IFITM-mediated restriction of virus production, identified the viral replication process that is subverted, and determined a means by which HIV-1 may partially overcome this effect. In addition, we find that restriction of viral production by IFITM proteins is clear contributor towards the inhibition of HIV-1 production seen in cells exposed to type I interferon, and have also shown that inhibition occurs in primary CD4^+^ T-cells at physiologically relevant levels of endogenous IFITM expression. From a mechanistic perspective we have shown that HIV-1 RNA is the target of this inhibition, though the viral determinant remains elusive. We therefore propose that inhibition of viral protein synthesis is a genuine antiviral function exerted by IFITM proteins that is distinct from previously described effects upon entry and infectivity.

Initial suggestions of a late-stage IFITM-mediated restriction have typically surrounded two common laboratory HIV-1 strains, NL4-3 and BH10 ^10,12,14,25,26^. However, for the putative anti-HIV restriction factor viperin/RSAD2, analysis demonstrated that the phenotype could not be replicated beyond the laboratory strain NL4-3 ^44^. Therefore our demonstration that IFITM proteins restrict production of multiple HIV-1, HIV-2 and SIVs implies the phenotype we observe is likely not an artefact. This is supported by our finding that African Green Monkey IFITM1 can exert a similar antiviral effect upon HIV-1 production as occurs with human IFITM1.

The general mechanism of restriction we outline is specific exclusion of HIV-1 mRNA from polysomes during IFITM expression. In HEK293T cells the effect principally occurs with IFITM1 and IFITM2, but less so for IFITM3 was generally consistent throughout our analysis. Yet inducible expression of IFITM3 in SupT1 cells showed an equivalent antiviral effect to IFITM1 and IFITM2, though, and this was also linked to viral protein loss and was independent of IFITM-driven effects on viral entry. We consider that the same translational blocks measured in polysome analysis of HEK293T cells drive the result seen in SupT1 cells and primary CD4^+^ T-cells. Indeed, we considered it very likely cell type specific differences would occur, as IFITM localization and thus antiviral activity are governed by both ubiquitylation and palmitoylation ^24,45^. Reasonably the degree to which IFITM-driven viral entry inhibition or viral translation inhibition occurs in particular cells is then influenced by the activity of these pathways, which are known to commonly vary. Equally, other cell specific factors influencing viral transcription (such as NFkB translocation) might also account for variation in antiviral activity between cell lines, as higher virus transcriptional activity may overcome inhibition. However, it is important to note that we were able to clearly confirm the inhibition of viral production in primary human CD4^+^ T-cells by IFITM2 and IFITM1-3, demonstrating that inhibition of viral production by IFITMs occurs in HIV-1 target cells at physiologically relevant levels of endogenously expressed IFITM proteins.

Why IFITM1 and IFITM2 should act similarly in many instances, when IFITM2 and IFITM3 have closer amino acid sequence identity and display greater overlap in cellular localization is not yet clear ^10,19,46,47^. IFITM-mediated antiviral effects upon viral entry are principally determined by cellular localization, and we anticipate the inhibition of protein synthesis to be similar ^45,46,48^. This is pertinent for differences we find between IFITM2 and IFITM3. Despite their high sequence similarity, they typically show distinct cellular localization.

The exclusion of viral transcripts from polysomes mainly seemed to mainly affect unspliced and singly-spliced transcripts. Yet despite a lack of significance, there was consistent but weaker loss of multiply spliced transcripts from polysome fractions, coupled with multiply-spliced transcript enrichment in low sucrose fractions. This is perhaps indicative of some degree of multiply-spliced transcript translational blockage that explains the loss of Nef protein seen throughout our experiments.

We saw no change in total unspliced mRNA levels in IFITM expressing cells yet there was modest depletion of singly-spliced and multiply-spliced mRNA during IFITM1 and IFITM2 expression. Though this overall depletion of total viral mRNA levels may partially account for some loss of Vpu and Nef production, polysome analysis additionally demonstrated clear exclusion of these transcript classes from translation. Why these overall mRNA levels should be reduced is of interest. One possibility is a transcriptional inhibition or pre-translational degradation, but degradation of mRNAs that have been inhibited in translation is thought to be common ^49^. For example, the antiviral protein ZAP, which degrades retroviral mRNA has also been shown to stall translation prior to mRNA degradation, illustrating the fluid link between translational stalling and mRNA stability ^50,51^.

Susceptibility to inhibition occurred at the level of viral mRNA and we so sought to identify a viral RNA susceptibility determinant. Our data showing *gag* codon optimization relieves the late-stage IFITM inhibition of HIV-1 supports similar findings ^26^. Singly-spliced and unspliced transcripts that bear the RRE were typically most affected by IFITMs proteins in our analyses. Yet investigation of proviral constructs in which the RRE was substituted for the CTE element were inconclusive, making it difficult to reconcile a role for the RRE in determining inhibition, despite the appealing observation that RRE bearing transcripts are typically most affected. Notably, RRE bearing transcripts also have a non-human codon bias and surrounding splice sites that may prove to influence restriction ^52–55^. As such further analysis to identify any HIV-1 RNA determinant of inhibition will then be of much value, though currently our further attempts to identify the viral RNA determinant have been unsuccessful. A recent study demonstrated that viral RNA CG dinucleotide frequency drives ZAP-mediated viral restriction, rather than a specific RNA structural determinants, thus the determinant for IFITM-mediated restriction of protein synthesis may be similarly obscure ^56^. However, we were able to exclude the broader possibility of general translational arrest via IFITM expression. Other aspects of the mechanism remain elusive also. IFITMs have no reported RNA binding domain and so would require either an RNA-binding partner to directly achieve inhibition, or would need to be involved in a pathway that could influence translation in this manner.

If IFITMs are cofactors in direct inhibition of viral RNA translation, it is not yet clear how a membrane protein can affect both soluble protein translation and membrane protein translation, this may ultimately depend the absolute strictness of translational site partitioning, a concept which is under renewed debate ^57^. Further, typically a proportion of IFITM proteins in cells are not palmitoylated, and show less distinct membrane localization, conceivably this fraction may be important in influencing translation ^45^. One unexplored possibility is that the capacity of IFITM proteins to disrupt membrane fluidity or other membrane physical characteristics to inhibition viral-cell membrane fusion ^17,58^ on the cell surface and in endosomal compartments similarly affects rough endoplasmic reticulum membranes, leading to a disruption of membrane bound translational complexes. Though how such a process could be specific to virus translation is unclear.

We demonstrate that the lentiviral protein Nef can help overcome the late-stage inhibition of viral production exerted by IFITM proteins. Though notably this rescue is not complete. Our data may help explain accounts of a Nef driven enhancement of HIV-1 production ^40,41^. The ability to overcome IFITM-mediated inhibition of virus production was apparent with all HIV-1, HIV-2 and SIV *nef* alleles tested. Though the underlying mechanism of ability of Nef to counter IFITMs is unclear, we saw no degradation of IFITMs in the context of viral protein expression. One possible explanation may be re-trafficking, wherein Nef acts as an adaptor protein to direct IFITMs away from their site of action on translation. This fits well with the sensitivity of IFITM-mediated antiviral function to changes in IFITM localization ^45,46,48^, but also fits with the ability of Nef to redirect a multitude of membrane proteins from their site of action ^38,39,59,60^. This may be consistent with our observation that Nef rescues virus particle production in supernatants during IFITM expression, but does not rescue viral protein production.

We also found that, in multiple cycle infections, the benefit of Nef expression was only seen early in culture but was latterly lost. Cell-free infection is prevalent in the early stages of culture, but infection shifts towards a cell-to-cell mode in later stages ^42^. It has been proposed that this represents a means of escape from immune factors in HIV-1; as the high efficiency, high multiplicity of infection associated with cell-to-cell transmission can overcome restriction ^61,62^. For example, with the antiviral restriction factor tetherin, antiviral effect is either severely weakened or absent during cell-to-cell transmission of HIV-1 ^31,63,64^.

Thus we conclude that IFITMs are antiviral factors that can target HIV-1 translation. For retroviruses this seems to be a recurring weak point in replication strategy that the innate immune system has frequently evolved to target. For example, PKR senses viral double stranded RNA and halts translation ^65^, while Schlaffen 11 inhibits viral protein production by disrupting tRNA synthesis ^66^, and finally the protein ZAP which induces viral mRNA degradation also stalls viral translation ^51^.

The model we put forward wherein IFITM expression specifically disrupts viral mRNA translation may be applicable to other targets. Conceptually, other self and non-self RNAs may be affected also. It is then of interest that a recent report found that LINE-1 retrotransposon mobility could be inhibited by IFITM1 expression ^67^, which for a retrotransposon cannot of course arise from viral entry effects. Future characterization will therefore identify the viral RNA determinant of susceptibility, and the breadth of antiviral and RNA regulatory response driven by IFITM proteins.

## Methods

### Plasmids

Human pQCXIP IFITM1-3 plasmids bearing an N-terminal FLAG were previously described ^58^. The viral construct pBR4-3-eGFP-Nef, encoding for wild type HIV-1 NL4-3, pBR4-3-IeGFPΔNef, HIV-1 NA7, HIV-1 NL4-3 plasmids carrying *nef* alleles of HIV-2 Ren or CBL and SIV AGM or MAC strains - all described in ^43^ - are kind gifts of Professor Frank Kirchhoff (University of Ulm). HIV-2 MCR and MCN plasmids ^68^ are kind gifts of Professor Áine McKnight (Queen Mary University of London). pSIV_agm_Tan-1, pSIV_cpz_Tan1.910 and pSIV_mac_1A11 are from NIH AIDS Reagent Program. Codon-optimized HIV-1 NL4-3 is encoded by pCNC-SYNGP ^33^ (Oxford BioMedica). Plasmids pCTEΔEnvΔRevΔRRE and pCTEΔEnvΔRev ^35^ are kind gifts of Professor Paul Bieniasz (Rockefeller University).

### Antibodies, cells and antiviral compounds

The following antibodies were used to detect IFITMs: human IFITM1 (clone 5B5E2, Proteintech), IFITM2 (clone 3D5F7, Proteintech) and human IFITM3 (clone EPR5242, Novus Biologicals). Anti-Flag (clone M2, Sigma) was used to detect FLAG-tagged IFITMs, Rev and GFP. Anti-HIV1 Nef (clone 3F2, ThermoFisher), rabbit polyclonal anti-Vpu (Abcam) and rabbit polyclonal anti-HIV1 Gag (Abcam) were used to detect viral proteins. As loading control, β-actin (clone AC74, Sigma) and GAPDH (Abcam) antibodies were used. Secondary antibodies were horseradish peroxidase conjugated goat anti-rabbit/mouse IgG. TZM-bl cells expressing shRNAs targeting IFITMs were previously described ^27^, as were SupT1 cells inducible for IFITM expression ^10^. Human Embryonic Kidney (HEK) epithelial 293T cells (HEK293T), C8166 cells, TZM-bl cells, and SupT1 cells were maintained in standard conditions, and were all originally procured from the NIH AIDS Reagents Program. Primary monocytes and CD4^+^ T cells were isolated from leukocyte cones (NHS Blood Transfusion service, St. George’s Hospital, London) using the human CD14^+^ and CD4^+^ T cell isolation kits, respectively, according to manufacturer instruction (Miltenyi Biotec). 0.5×10^6^/ml CD14^+^ monocytes were differentiated into macrophages with 100ng/ml GMCSF (Peprotech) at 2-3 days interval until day 7. CD4^+^ T cells were activated at 1×10^6^/ml with T cell activator CD3/CD28 Dynabeads (ThermoFisher), at a bead-to-cell-ratio of 1:1, in complete medium (RPMI 1640 medium supplemented with 10% heat-inactivated fetal bovine serum, 2mM L-glutamine, 100U/ml streptomycin, 100U/ml penicillin, all from ThermoFisher) and 30U/ml IL-2 (Peprotech) for 48 hours. AMD3100 and Maraviroc were obtained via the NIBSC AIDS Reagent Program.

### Transfections

HEK293T cells were plated at 2×10^4^/cm^2^ in 48-well plates (for measurements of virus production), 6-well plates (for measurements of whole cell viral transcripts by qPCR), 10cm dishes (for virus production, polysome analysis and immunoprecipitation, all from Nunc), 48 hours before transfection. Plasmid DNA (total quantity of 1μg/well in 48-well plates, 3μg/well in 6-well plates and 5μg/well in 10cm dishes) was diluted in OptiMEM (ThermoFisher) at 10% volume of total cell culture medium. Equal quantity of different plasmid DNA was added in co-transfection. Linear polyethylenimine (L-PEI) was added at a w/w ratio of 5:1 into the diluted DNA. The transfection mixture was incubated at room temperature for 10 minutes before added to cells.

For siRNA transfection experiments, HEK293T cells were plated at 1×10^4^/cm^2^ 24 hours prior to transfection in 48-well plates. siRNAs at a final concentration of 100nM was added to a total of 100μl OptiMEM, incubated for 5 minutes at room temperature and then 1μl DharmaFECT 1 (Dharmacon) was added to diluted DNA. siRNA-transfection mix was incubated for 20 minutes at room temperature and then added to cells. Expression vectors of IFITMs and HIV-1 proviral DNA were transfected then 48 hours post-transfection.

TZM-bl cells were plated at 6×10^4^/cm^2^ in 48-well plates 24 hours before transfection. In a 48-well format, 1μg/well plasmid DNA and 6μl Lipofectamine 2000 (ThermoFisher) were diluted in OptiMEM, incubated for 10 minutes at room temperature and then added to cells. Medium was replaced with medium containing 5μM AMD3100 and 100IU/ml IFN-β (Peprotech) 4 hours post-transfection.

### Intracellular staining and flow cytometry

IFITM expression in IFNβ-treated human MDMs, CD4^+^ T cells, transfected HEK293T cells and doxycycline-treated SupT1 cells was detected by fixing cells in 4% (w/v) paraformaldehyde (Sigma) in PBS for 15 minutes at room temperature, followed by permeabilization in 0.2% (v/v) Triton X-100 (Sigma) for 20 minutes at room temperature. Cells were then blocked in 5% BSA in PBS for 1 hour at room temperature followed by incubation with 1μg/ml IFITM-specific monoclonal antibodies diluted in staining buffer (2% BSA/0.1% Triton X-100/PBS) overnight at 4°C. Cells were washed 3 times with 2% BSA in PBS and labelled with secondary antibodies, human IFITM1, 2 and 3 antibodies were detected by 0.5μg/ml goat anti-mouse IgG2a Alexa Fluor 647, goat anti-mouse IgG1 Alexa Fluor 488 and goat anti-rabbit Alexa Fluor 350, respectively, in staining buffer for 1 hour at room temperature. Labelled cells were washed 3 times in 2% BSA in PBS and analyzed on LSR II flow cytometer (BD). Data was analyzed on FlowJo (BD).

### Infection

HIV-1 virus was produced from transfected HEK293T cells. SupT1 cells were spinoculated with the indicated concentrations of inoculum for 2 hours at 37 C at 1000x *g* in 96-well U-bottom plates (Nunc) followed by incubation at 37 C for 1 hour. Infected cells were then washed 3 times with phosphate-buffered saline (PBS, Sigma) and re-suspended in medium with 1μg/ml doxycycline (Sigma) and 5μM AMD3100. C8166 cells were infected with the same protocol without AMD3100. Supernatant was harvested at the indicated time-points after centrifugation at 500x *g* for 5 minutes at room temperature.

### Transduction and infection of primary human CD4^+^ T cells

In the presence of CD3/CD28 T-cell activator beads, 5×10^5^/ml activated CD4^+^ T cells were transduced with IFITM-targeting or scrambled shRNA lentivirus with p24 at 100ng/ml for 2 hours at 1000 x*g*, 37°C and then for 1 hour incubation. Infected cells were washed three times with PBS, re-suspended in complete medium with 30U/ml IL-2 and then incubated at 37°C for 48 hours. Prior to HIV infection, T cell activator beads were removed from transduced CD4^+^ T cells. Transduced CD4^+^T cells were then incubated with medium alone, 5μM Maraviroc or 5μM AMD3100 for 2 hours at 37°C. Untreated cells and drug-treated cells were then infected with 100ng/ml p24 of NL4-3 or 89.6 virus, respectively, at 37°C, 1000 x*g* for 2 hours and then incubated for 1 hour. Infected cells were then washed three times with PBS and re-suspended in complete medium containing 30U/ml IL-2, 5μM Maraviroc and 5μM AMD3100. Aliquots of infected cells were treated with 0.25% Trypsin/EDTA (ThermoFisher) for 15 minutes at 37°C, washed three times with PBS, fixed in 2% paraformaldehyde, permeabilized in 0.2% Triton X-100/PBS and stained with RD-1 conjugated monoclonal mouse anti-HIV-1-p24 (clone KC57, Beckman Coulter) for flow cytometry analysis of internalized virus. Level of viral output in supernatant was measured by p24 ELISA 72 hours post-infection.

### p24 ELISA

ELISA plates (Nunc) were pre-coated with 5μg/ml sheep anti-HIV-1 p24 antibody (Aalto Bio Reagents) at 4 C overnight. Supernatant of transfected HEK293T, TZM-bl cells or infected cells was treated with 1% Empigen BB (Sigma) for 30 minutes at 56°C, then plated at 1:10 dilution in Tris-buffered saline (TBS) on anti-p24-coated plates and incubated for 3 hours at room temperature. Alkaline phosphatase-conjugated mouse anti-HIV-1 p24 monoclonal antibody (Aalto Bio Reagents) diluted in 20% sheep serum, 0.05% v/v Tween-20, TBS (all from Sigma) was then added and incubated for 1 hour at room temperature. Plates were washed 4 times with 0.01% v/v Tween-20 in PBS and twice with ELISA Light washing buffer (ThermoFisher). CSPD substrate with Sapphire II enhancer (ThermoFisher) was added and incubated for 30 minutes at room temperature before chemiluminiscence was read by a plate reader.

### Reverse transcriptase (RT) activity and luciferase activity assays

Colorimetric reverse transcriptase activity assay kit (Roche) was used to determine reverse transcriptase activity of HIV-1, HIV-2 and SIV in supernatants of transfected HEK293T cells. Manufacturer instructions was followed except that supernatant was first treated directly with lysis buffer (50mM Tris pH7.8, 80mM potassium chloride, 2.5mM DTT, 750μM EDTA and 0.5% Triton X-100, all from Sigma) for 30 minutes at room temperature and then incubated in streptavidin-coated microplates for 15 hours at 37 C.

Luciferase activity of infected TZM/bl cells was analyzed with Bright-Glo luciferase activity kit following manufacturer instructions (Promega).

### Polysome profiling

Polysome analysis was performed with both manual fractionation and RNA analysis ^69^. Transfected HEK293T cells were incubated with 100μg/ml cycloheximide (Sigma) for 15 minutes for 37°C and then washed with ice-cold PBS with 100μg/ml cycloheximide. Cells were then lysed in polysome buffer, 10mM Tris pH8, 140mM NaCl, 1.5mM MgCl_2_, 0.5% v/v NP40, 100μg/ml cycloheximide, protease inhibitor cocktail (all from Sigma) and 800U/ml RNase OUT (ThermoFisher), for 10 minutes on ice. Cell lysate was centrifuged at 10, 000x *g* for 1 minute at 4°C and supernatant was then adjusted to 200μg/ml cycloheximide and 700μg/ml heparin (Sigma). Following centrifugation at 12, 000x *g* for 10 minutes at 4°C, an aliquot of supernatant was taken as input and the rest was layered onto a 10% to 50% sucrose gradient prepared using polysomal buffer. Gradients were then ultracentrifuged at 300,000x *g* for 16 hours at 4°C (Sorvall). After centrifugation, 20 550μl fractions were collected from the top of the gradients for immunoblotting and quantitative PCR analyses. Level of RNA in fractions was measured by absorbance at 254 nm (Nanodrop) and RNA was precipitated with 5x volume of absolute ethanol (Sigma) overnight at −20°C.

### Quantitative PCR

Total RNA of transfected HEK293T cells and precipitated RNA from polysome profiling fractions were purified with RNeasy mini kit and on-column DNA digestion with RNase-free DNase kit (both from Qiagen) following manufacturer instructions. 3ng of total RNA per sample was analyzed with Superscript III Platinum One-Step qRT-PCR kit with ROX (for unspliced and multiply spliced viral transcripts, ThermoFisher) or QuantiTect SYBR Green PCR kit (for singly-spliced viral transcript, Qiagen) and ABI 7500 Real Time PCR system (Applied Biosystems). Cycling conditions were 50°C C for 15 minutes, 95°C C for 8 minutes, then cycling of 95°C for 15 s and 60°C for 30 s. Reactions carried out in the absence of reverse transcriptase (Platinum *Taq* only) confirmed the absence of DNA contamination. The samples were quantified against cloned standards.

Primers used to amplify unspliced (Forward, 5’-CCGTCTGTTGTGTGACTCTGG-3’, reverse, 5’-GAGTCCTGCGTCGAGAGATCT-3’), multiply-spliced (Forward, 5’-CAGACTCATCAAGCTTCTCTATCAA-3’, reverse, 5’-CTATTCCTTCGGGCCTGTC-3’) and singly-spliced (Forward, 5’-TAATCGGCCGAACAGGGACTTGAAAGCGAAAG-3’, reverse, 5’-CCCATCTCCACAAGTGCTGATACTTC-3’) viral transcripts are described in ^70^, ^71^, and ^72^, respectively. Oligonucleotide probes are labelled with 5’-FAM and 3’-TAMRA, (unspliced, 5’-TCTAGCAGTGGCGCCCGAACAGG-3’ and multiply-spliced, 5’-AACCCACCTCCCAATCCCGAGG-3’, all from ThermoFisher). Cellular GAPDH (glyceraldehyde-3-phosphate dehydrogenase) mRNA was additionally amplified as a loading control with primers (Forward, 5’-AGGTCGGAGTCAACGG ATTTGG-3’, reverse, 5’-GATGGCAACAATATCCACTTTACCA-3’) and probe (5’-TCTTATTGGGCGCCTGGTCAC-3’, as described in ^71^.

### Immunoblotting

Cells were washed once with PBS and then lyzed in radioimmunoprecipitation buffer (RIPA, containing 20mM Tris pH7.5, 150mM NaCl, 1mM EDTA, 1mM EGTA, 1% NP40, 1% sodium deoxycholate, 250μM sodium pyrophosphate, 1mM β-glycerophosphate, 1mM sodium vanadate and protease inhibitor cocktail, all from Sigma) for 30 minutes at 4°C. Lysate was then centrifuged for 10, 000x *g* for 10 minutes at 4°C. Protein concentration of supernatant was determined by bicinchoninic acid (BCA) assay (ThermoFisher). 10μg of protein per sample was analyzed by immunoblotting, developed with ECL Prime reagents (GE Healthcare Life Sciences) and captured on CL-XPosure films (ThermoFisher) or ChemiDoc MP system (Bio-Rad). Intensity of immunobands was analyzed by ImageJ (on X-ray films) or ImageLab (Bio-Rad).

### Statistical analysis

Levels of viral output in p24 ELISA and RT assay were normalized to vector-transfected or scrambled shRNA-transduced control and expressed as the level of virus production unless otherwise indicated. Statistical analysis was performed with Graphpad Prism 5. Data shows mean + standard error of mean from a minimum of 3 independent experiments.

## Acknowledgements

We thank Áine McKnight for helpful discussions regarding this work and Greg Towers for useful comments on this manuscript. RDS was supported as a Barts and The London School of Medicine and Dentistry Early Career Research Fellow. We thank Frank Kirchhoff, Paul Bieniasz, and both the NIH and NIBSC AIDS reagent programmes for the provision of reagents.

## Author Contributions

Designed the study: WYL, CL, RDS. Performed the experiments: WYL, RDS. Analyzed the data: WYL, RDS. Wrote the manuscript: WYL, RDS. Contributed reagents: CL.

## Supporting Information Figure legends

**Supporting Fig 1. IFITMs inhibit HIV-1, HIV-2 and SIV viral output.** Levels of virus production of HEK293T cells transfected with 0.5μg expression vectors of IFITMs and 0.5μ**g (A)** HIV-2 strains or **(B)** SIVs (AGM –African Green Monkey, CPZ – Chimpanzee, MAC – Macaque) proviral DNA were measured by RT activity assay and normalized. **(C)** Levels of virus production of HEK293T cells transfected with 0.5μg expression vectors of African Green Monkey IFITM1 and 0.5μg proviral DNA of HIV-1 were measured by p24 ELISA and normalized. Data shows mean+SEM of 3 independent experiments. Differences were analyzed with Student’s t-test and * indicates p<0.05.

**Supporting Fig 2. AMD3100 blocks HIV-1 NL4-3 entry in SupT1 cells.** 1×10^6^/ml SupT1 cells were treated with 5μM AMD3100 or DMSO for 2 hours at 37°C and then left untreated or infected with HIV-1 NL4-3 with p24 concentration of 100ng/ml for 2 hours by spinoculation and 1 hour incubation at 37°C. Cells were then washed 3 times with PBS. Levels of virus production were measured by p24 ELISA 48 hours post-transfection. Mean fluorescence intensity of IFITMs in **(B)** human monocyte-derived macrophages (MDMs, day 7) and **(C)** human CD4^+^ T cells treated with indicated concentrations of IFNβ for 24 hours; **(D)** HEK293T cells transfected with 0.5μg IFITM-encoding plasmids and 0.5μg HIV-1 proviral DNA for 48 hours and **(E)** SupT1 cells treated with 1μg/ml doxycycline for 48 hours to induce expression of the indicated IFITMs, was measured by intracellular staining of IFITMs with monoclonal antibodies and flow cytometry. Data shows mean+SEM of 3 independent experiments.

**Supporting Fig 3. Substituting the MPMV CTE for the HIV-1 RRE does not affect IFITM-mediated inhibition of HIV-1 production.** HEK293T cells were transfected with expression vectors for IFITMs and HIV-1 NL4-3 DNA (wild type [*i.e.* RRE bearing], MPMV CTE only or CTE+RRE). Level of viral production was measured by p24 ELISA 48 hours post-transfection and normalized. Data shows mean+SEM of 3 independent experiments. Differences were analyzed with Student’s t-test and * indicates p<0.05.

**Supporting Fig 4. (A) HIV-1 Nef does not overcome IFITM-mediated inhibition of early viral replication steps.** C8166 CD4^+^ T-cells were transduced to constitutively overexpress IFITM1-3. Cells were then infected with the wild type pBR4-3-eGFP-Nef virus or pBR4-3-eGFP-ΔNef virus. Cells were measured for GFP expression 48h post-infection by FACS to determine infection rate. Data shows fold-change in restriction of GFP infection rates during IFITM expression relative to empty vector control cells. **(B)** Levels of p55 and p24 in immunoblotting of Fig. 7B were quantified by densitometry and normalized to β-actin. Data shows mean + SEM of the ratio of viral proteins to β-actin of 3 independent experiments and differences were assessed by Student’s t-test, * denotes p<0.05. **(C) Increasing Nef levels rescues HIV-1 protein production during IFITM expression.** HEK293T cells were transfected with ΔNef HIV-1 NL4-3 proviral DNA, expression vectors for IFITMs and an increasing proportion of HIV-1 Nef-encoding vector versus empty vector in a fixed total quantity. Levels of viral proteins and IFITM-FLAG expression was analyzed by immunoblotting and densitometry. Data show mean + SEM of 3 independent experiments and differences were assessed by Student’s t-test, * denotes p<0.05.

